# Ability of mycobacterial pathogens to establish a persistent infection is lost by deletion of a single gene, *rel*, regulator of the stringent response

**DOI:** 10.1101/2024.01.10.575042

**Authors:** Asmaa H. Mahmoud, Gaber S. Abdellrazeq, Lindsay M. Fry, David A. Schneider, Sarah Attreed, Leeanna Burton, Neha Sangewar, Waithaka Mwangi, Cleverson deSouza, Victoria Hulubei, William C. Davis, Kun Taek Park

## Abstract

Studies in a mouse model revealed *Mycobacterium tuberculosis* (*Mtb*) with a deletion of *rel*, regulator of the stringent response, could not establish a persistent infection. Studies in cattle with a *Mycobacterium. a. paratuberculosis rel* deletion mutant revealed inability to establish a persistent infection was associated with development of CD8 cytotoxic T cells (CTL) that kill intracellular bacteria. Further comparative studies ex vivo with *Mbv* Calmette-Guérin (BCG) and a BCG *rel* deletion mutant revealed no clear difference in development of CTL in vitro. As reported, a study of the recall response was conducted with cattle vaccinated with either BCG or with BCG*rel,* to determine if information could be obtained that would show how gene products under control of *rel* interfere with the CTL response to mycobacterial pathogens in vivo. The study revealed the CTL response elicited by vaccination with BCG was impaired, in comparison with the response elicited by BCG*rel*. Comparative analysis of the recall response ex vivo revealed the functional impairment was not associated with the timing of appearance of the recall response, expression of IFN-γ, TNF-α, IL-17, or IL-22, or molecules that mediate intracellular killing. Further studies are needed to determine how CD8 CTL functional activity is modulated in vivo by gene products regulated by *rel*.

## 1. Introduction

Extensive studies have been conducted since the discovery that *Mycobacterium tuberculosis* (*Mtb*), *M. bovis* (*Mbv*), and *M. avium subsp. paratuberculosis* (*Map*) are the causative agents of tuberculosis and paratuberculosis (1, 2). The studies have revealed *Mtb*, *Mbv*, and *Map* are members of a much larger group of mycobacteria that cause disease in humans and animals (reviewed in (3–6)). This has necessitated a change of research focus to a more global approach to understanding immunopathogenesis of mycobacterial infections and development of vaccines. A seminal observation made by investigators studying the stringent response has opened up an additional new opportunity to reveal the mechanisms used by mycobacteria to evade immune clearance and establish of persistent infection (7). Lack of understanding how mycobacteria evade immune clearance has been the major impediment to developing effective vaccines for mycobacterial pathogens. The search for gene products that prevent establishment of a persistent infection has not been successful. Studies with mycobacterial deletion mutants and candidate peptide vaccines have elicited immune responses that modulate the immune response but none of them have elicited a response that prevents establishment of a persistent infection (8–10). The mechanisms used by mycobacterial pathogens to evade immune clearance seemed to be intractable. However, Dahl et al. have identified a gene that does affect the capacity of *Mtb* to establish a persistent infection. They observed, in a mouse model, that deletion of the gene *rel*, regulator of the stringent response, abrogates the capacity of *Mtb* to establish a persistent infection (7). They noted that the mutant was able to establish an infection with development of granuloma lesions in the lung and spleen, identical to those that develop following infection with wild type *Mtb*. In contrast to infection with *Mtb* the granulomas resolved by 15 weeks post infection with clearance of the mutant from all tissues. The interpretation at that time was that inability to survive was associated with loss of ability to adapt to survival in a nutrient limited environment, owing to the absence of signaling by genes under the control of *rel*.

The significance of the studies by Dahl et al. has not attracted very much attention, because the interpretation of the observations was that nutrient starvation was the reason why the *rel* deletion mutant was unable to survive in the in vivo environment of the vertebrate host. However, it did attract our attention. The ability of the mutant to establish an infection and persist long enough to induce the initial stages of pathology before clearance suggested that clearance was associated with development of an immune response that cleared the infection. It also suggested that products encoded by genes under the regulation of *rel*, were involved in preventing immune clearance. As reported in our initial studies, we were interested in determining whether the presence of *rel* was a universal requirement for mycobacterial pathogens to establish a persistent infection. We chose to use cattle as a model species to explore this possibility. We selected *Map* as another lineage of pathogenic mycobacteria to determine if *rel* is essential for survival in vivo. To determine if the deletion of *rel* is unique we deleted the orthologue of *rel* in *Map* (11). For comparison we developed a mutation in *pknG,* a gene known to affect survival of *Mbv* and BCG in macrophages (11) (12). A study using cattle and goats revealed the *pknG* was able to establish a persistent infection. In contrast the *Maprel* mutant was cleared from all tissues as detected by culture for bacteria and PCR (13). Methods were developed and used to study the immune response to mycobacteria ex vivo in real time, starting with *Maprel*. Cattle were vaccinated with *Maprel* or *Map*. PBMC were cultured with *Map* or *Maprel*. A CD4 CD8 T cell recall response was demonstrated with use of flow cytometry (13). Further studies were conducted to study the recall response elicited with antigen presenting cells (APC) (14). Analysis of the recall response using APC primed with either *Maprel* or a candidate vaccine membrane protein (MMP) revealed both stimulants elicited a CD4 CD8 proliferative recall response (14). Further analysis of the primary immune response using APC primed with *Maprel* or MMP to stimulate peripheral mononuclear cells (PBMC) demonstrated stimulation elicited a proliferative response in memory CD4 CD8 T cells. The CD8 T cells developed into cytotoxic T cells with ability to kill intracellular bacteria (15). No response was detected in γδ T cells or NK cells.

To extend the studies and obtain more information on the role of *rel* in establishment of a persistent infection, we developed a *rel* deletion mutant in the *Mbv* Calmette-Guérin (BCG) vaccine. As reported in our initial report, we were interested in determining if a difference could be detected in the recall response to BCG and BCG*rel* ex vivo in tissue culture. Multiple studies with BCG have shown that, although virulence is reduced, BCG can still establish a persistent infection. This was demonstrated in early studies reported by Lurie in 1934 (16) and more recent studies reviewed in (8, 17, 18). It could be inferred from the *Mtb* studies by Dahl et al, that deletion of *rel* in BCG would have the same effect as deletion in *Mtb*. The potential of a *rel* deletion mutant for improving the efficacy of BCG was the rationale for switching from *Map* to BCG to continue the studies. The development of assays to study the immune response in real time provided an opportunity to compare the immune response ex vivo using a tissue culture platform (19). Comparison of the immune response following stimulation with APC primed with BCG*rel* or with BCG revealed no significant difference in the proliferative response. No significant difference was observed in expression of 3 cytokines (IFN-γ, TNF-α, and IL-17) that modulate the immune response. Likewise, no difference was observed in the content of 3 molecules, (perforin, Granzyme B, granulysin) known to play a role in intracellular killing of bacteria. Of special interest, no difference was observed in the functional activity of CD8 T cells that develop following stimulation with APC primed with BCG*rel* or BCG ex vivo. Both methods of stimulation led to development of CD8 T cells with equal ability to kill bacteria present in infected target cells (19).

For the ex vivo studies, there was limited opportunity for mycobacteria to express products encoded by genes under the regulation of *rel*. As described in the present follow up report, studies were conducted to compare the immune response to BCG and BCG*rel* in vivo through analysis of the recall response ex vivo. In the in vivo environment, products secreted by mycobacteria could modulate Ag processing and signaling through APC leading to changes in the phenotype and/or function of memory CD4 and CD8 T cells at the time of Ag presentation or later. We compared 1) phenotype of memory CD4 and CD8 T cells proliferating in response to stimulation with APC primed with BCG*rel* or BCG, 2) expression of regulatory cytokines IFN-γ, TNF-α, IL-17, and IL-22, 3) content of perforin, Granzyme B, and granulysin in CD4 and CD8 memory T cells from cattle vaccinated with BCG*rel* or with BCG. We also compared the functionality of the memory CD4 and CD8 T cells after stimulation with APC primed with live BCG*rel* or with BCG.

## 2.. Materials and methods

### 2.1. Animals

Fifteen naïve Holstein steers were obtained from the *Map* free Washington State University (WSU) Knott Dairy. Three of the naïve steers, used to obtain data on the ex vivo primary recall immune response in our initial studies, were also used in the present study. Two of the steers were vaccinated with either BCG or BCG*rel.* The third steer was kept as an unvaccinated control. Two additional steers were obtained and used to verify the earliest time point a recall response could be detected following vaccination. Finally, ten steers (born in January 2020) were used for the main study. The steers were grouped as follows: One group of 4 steers was vaccinated with a live Danish BCG strain. A second group of 4 steers was vaccinated with live BCG*rel*. The third group of 2 steers was used as controls. Immunization consisted of a single intradermal injection of 10^9^ live BCG or BCG*rel*. The steers were kept in an open feedlot and used as a source of blood to conduct the studies. Steers were maintained by the WSU animal care staff. All procedures involving the use of animals were approved by the WSU Institutional Animal Care and Use Committee (ASAF 6542).

### 2.2. Bacterial strains and culture conditions

The *Mbv* Danish BCG strain, obtained from NADC (Ames, IA, USA), was used for BCG immunization and to develop a BCG*rel* deletion mutant [13, 19]. The Danish strain is currently used as a vaccine strain in humans and for comparative studies of the immune response to pathogenic *Mbv* in cattle. The seminal novel method of gene deletion developed by Bardarov et al. was adapted and used to develop a BCG*rel* deletion mutant (11, 20). Bacteria were cultured as described in (15). Prior to immunization, bacteria were pelleted via centrifugation at 4500 × g for five minutes, the supernatants were discarded. The pellets were resuspended in PBS (pH 7.4) and disaggregated. The optical density at 600 nm (OD600) was used to estimate the bacterial number and used to dilute the bacteria to a multiplicity of infection (MOI) needed to conduct the studies described below.

### 2.3. Blood processing and cell culture

Blood samples (∼ 200 mL) were collected from each steer via jugular venipuncture in weeks 2, 4, 6, and 8 post-vaccination (PV) for each biological experiment. PBMC were isolated from buffy coats using density gradient centrifugation and then by overlaying buffy coats, diluted two-fold in phosphate-buffered saline/Acid Citrate Dextrose 10%, pH 7.4 (PBS/ACD), over Ficoll-Paque™ PREMIUM, density1.077/ml (Cytiva UT). The interface containing PBMCs was collected and washed. Residual red blood cells were lysed using tris-buffered ammonium chloride (0.83% NH4Cl, 0.1% KHCO3, 0.01M EDTA pH 7.2). PBMC were washed three times with PBS/ACD then resuspended in complete culture medium (cRPMI) RPMI-1640 medium with GlutaMAX™ (Life Technologies, CA) supplemented with 10% bovine calf serum (HyClone, USA) and 50 μM β-mercaptoethanol. PBMC were counted and plated in 6-well tissue culture plates (2 × 10^6^/mL in 5 mL of warm cRPMI). Live BCG and BCG*rel* were added to duplicate wells (5 × 10^6^/ well, MOI 0.5:1). One well remained unstimulated, and one well was stimulated with 5 ug/ml Concanavalin A (conA), each to serve respectively as negative and positive controls. All cultures were incubated at 37°C / 5% CO2 for 6 days.

### 2.4. Flow cytometric analysis of the recall responses of BCG and BCGrel-stimulated PBMC from vaccinated and unvaccinated steers

To determine the recall response, blood was collected on weeks 2, 4, and 6 PV. For each time point, PBMC were cultured as described above and incubated for 6 days. On day 6, cultured PBMC were collected, washed twice with PBS, counted, and plated in 96 well polystyrene V-shape bottom microplates (10^6^ cells/well). The cells were washed twice with first wash buffer (FWB: PBS containing 0.01% w/v sodium azide, 0.02% v/v horse serum, and 10% v/v acid citrate dextrose) and processed for flow cytometry (FC), labeling as previously described (14, 21). In brief, the cells were incubated with mAbs listed in Table 1 (1 μg of mAb/10^6^ cells) for 15 min in the dark on ice. Cells were then washed twice using FWB and resuspended in 100 μL of fluorochrome-conjugated goat anti-mouse isotype-specific secondary mAbs. Appropriate isotype controls were used in all experiments. Cells were incubated for 15 min in the dark on ice, then washed twice using a second wash buffer SWB (the same as FWB but without horse serum). After the final wash, cells were resuspended in PBS buffered formaldehyde (2%) and stored at 4°C until examined by FC.

FC analysis of the proliferative response of CD4 and CD8 T cells to BCG or to BCG*rel* stimulation was performed as previously described (19). Data were acquired using the BD FACS Calibur (BD, USA). A sequential gating strategy was used to accurately define the target T cell subsets as shown in Fig. 1.

**Figure 1.**
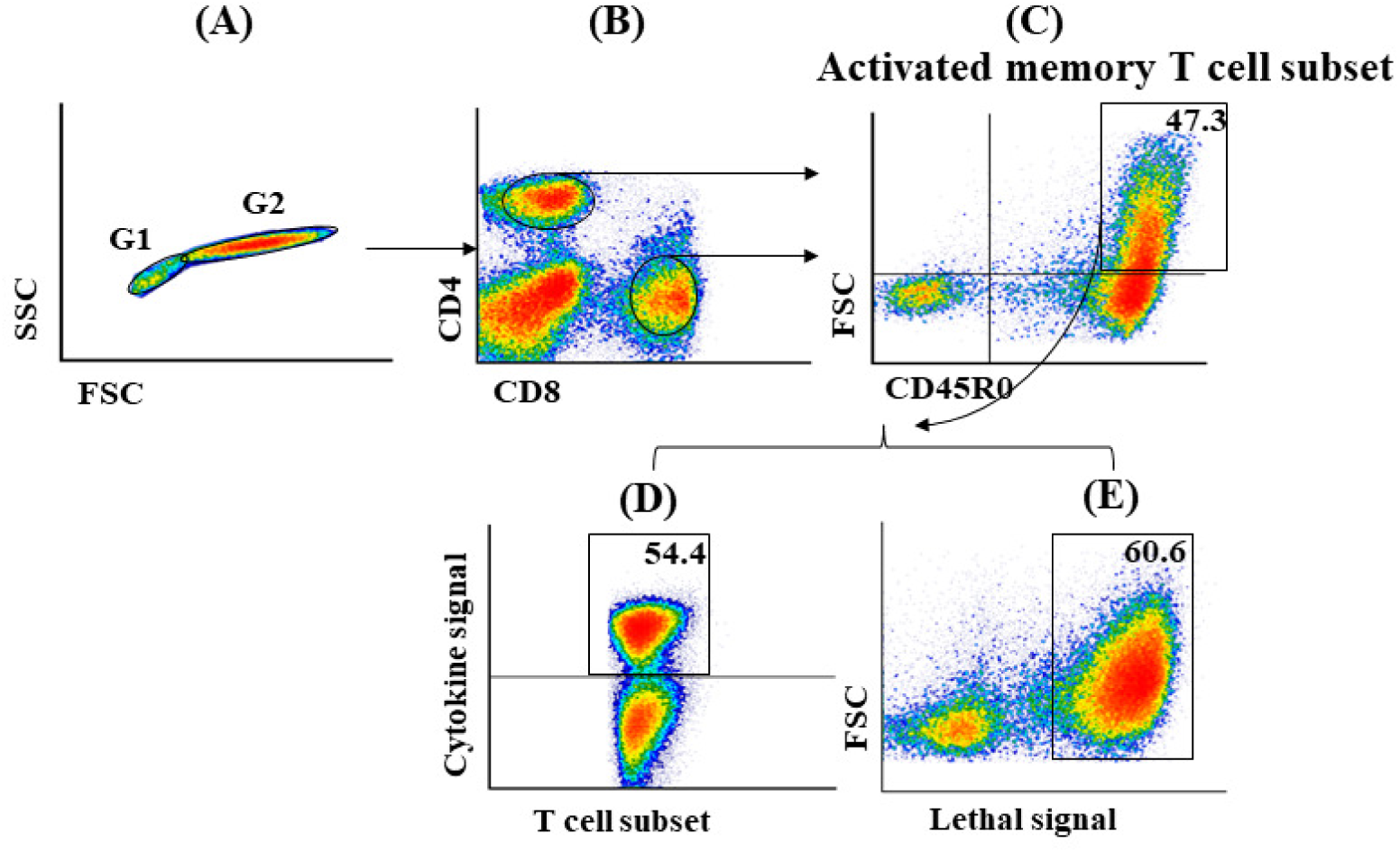
Flow diagram illustrating gating strategy used to isolate CD4 and CD8 lymphocytes for analysis. CD4 and CD8 T cells were labeled with mAbs specific for CD4 and CD8, the memory T cell marker CD45R0, cytokines (IFN-γ, TNF-α, IL-17, IL-22), and mAbs specific for molecules that provide the lethal signals that kill intracellular bacteria (perforin, GranzB, and granulysin). Different combinations of second step goat isotype specific antibodies conjugated with different fluorochromes were used to distinguish the mAbs specific for the different molecules. (A) Two electronic gates were placed on lymphocytes to exclude non-specific signal, SSC vs FSC. Signal was visualized as density plots. Gate 1 was placed on small non-activated lymphocytes. Gate 2 was placed on activated lymphocytes. (B) Gates 3 and 4 were used to isolate CD4 and CD8 T cells for further analysis. (C) CD4 and CD8 CD45R0 positive memory cells were visualized in Gate 5 FSC vs CD45R0 to electronically isolate the respective activated memory cells for further analysis. CD4 and CD8 positive activated memory cells were visualized in 2 additional gates (D) to distinguish CD45R0 positive memory cells that expressed the respective cytokines and (E) to distinguish CD45R0 positive memory cells that expressed respective lethal signal molecules.

### 2.5. Cytokine and lethal signals of BCG and BCGrel stimulated PBMC from vaccinated and unvaccinated control steers

To determine the FC profiles, blood was collected at weeks 2, 4, and 6 PV to analyze cytokine production and in week 8 PV to detect the lethal signals mediated by perforin, GranzB, and granulysin. PBMC were cultured as described above and incubated for 6 days. On day 6, Brefeldin A (BFA; 3 μg/mL, BD Biosciences) was added for the last 4 hours of incubation to block the secretion of proteins. The cultured PBMCs were harvested. Surface staining was carried out as described above. Then intracellular staining was carried out as described previously (19, 22). In brief, after surface staining for T cell subsets, cells were fixed and permeabilized using BD CytoFix/CytoPerm Fixation Permeabilization solution (BD Biosciences, USA) for 20 min on ice. Cells were washed twice with 1X perm/wash buffer (BD Biosciences, USA) and resuspended in perm/wash buffer containing 1 μg of mAb conjugated with Zenon 647/ 488 IgG1 (ThermoFisher Scientific, USA) for 30 min in the dark on ice. The mAbs used in the study are listed in Table 1. Different combinations of Zenon goat anti-IgG1 bodies AlexaFluor 647 or 488 (ThermoFisher Scientific, USA) conjugated with anti IFN-γ, TNF-α, IL-17A, IL-22 perforin, Granzyme B, or granulysin were used to detect the expression of cytokines and lethal signals on memory CD4 and CD8 T cells. In some experiments, cells were labeled with anti-IFN-γ conjugated with Alexa Fluor 647 and with either anti-TNF-α, or -IL-17A, or -IL-22 conjugated with Alexa Fluor 488, to determine whether the cytokines were differentially expressed in subsets of CD4 and CD8 T cells. Intracellular labeling of controls was included (without the primary antibodies) to demonstrate the specific binding of antibodies. After the cells were incubated with conjugated mAbs for ∼ 30 min in the dark on ice, the cells were washed twice with 1X perm/wash buffer, re-suspended in PBS-buffered formaldehyde (2%), and examined immediately after labeling. Data were acquired and analyzed as described above. Approximately 5 × 10^5^ events were collected for each sample. The unstimulated, BCG, and BCG*rel* and stimulated cultures from unvaccinated and vaccinated steers were analyzed for the expression of the cytokines and lethal signals.

#### 2.5.1. Culture and processing of PBMC for determination of bacterial viability

Two identical sets of PBMC cultures were prepared in duplicate as described above. One set of PBMC were cultured with or without the stimulant (with BCG or BCG*rel*) for 6 days for generation of effector CTL cells. Another set of PBMC without stimulant were cultured to generate moMacs. This set was incubated overnight at 37°C / 5% CO2 to allow adherence of the monocytes. The majority of the nonadherent cells were removed by washing and replacing the medium the following day. This washing step was repeated on day 3. The culture sets were incubated for a total of six days to generate moMac cells from each steer.

On day 6, moMac were infected with BCG at an MOI of 10:1 (2 × 10^7^ BCG to ∼2 × 10^6^ moMac/well) to generate infected target cells as described in (15). Unstimulated, control PBMC or effector/CTL cells (referred to as E/CTL in the text) developed by stimulation with BCG or BCG*rel*) were collected, washed twice with warm PBS, resuspended in 5 ml of antibiotic-free RPMI at 2 × 10^6^ cells/mL, and added to the preparations of BCG-infected moMac in duplicate. The cultures were incubated for 24 hours at 37°C / 5% CO2.

On the following day, supernatants containing non-adherent cells were collected into new tubes and centrifuged at 20,000 × g for 10 min to pellet cells and bacteria. Supernatants were discarded and the pellets collected and recombined with corresponding adherent cells. The combined cell preparations were then permeabilized with a solution of saponin (0.5% w/v in PBS 10x, Thermo Fisher Scientific, USA) for 15 minutes at 37°C, to release intracellular bacteria. All cell lysates were collected and transferred into new 2 mL-tubes and centrifuged at 20,000 × g for 10 min. Pellets were washed and re-suspended in 500 μl of DNase-free H2O as described, then stored at - 20°C until use.

Another set of unstimulated PBMC and moMac were prepared and used as live and dead mixtures of bacteria to serve as controls (15, 23). In brief, six wells of unstimulated PBMC and moMac were prepared as described above. A set of controls were prepared from known mixtures of live and dead BCG as follows: Aliquots of BCG mixed in three ratios, 100% live, 50% live/50% killed, and 100% killed, were prepared in duplicate to obtain 2 × 10^7^ total BCG in each aliquot, then added to the cultures of moMac at a MOI of 10 and incubated for 3 hours. The cultures were washed to remove extracellular bacteria. To each well were added 10^7^ unstimulated PBMC and 2 mL diluted saponin solution, which was incubated at 37°C for 15 minutes to release intracellular BCG. The further steps were carried out as described above and lysates were stored at -20°C until used.

#### 2.5.2. PMA treatment

All bacteria-cell lysates were treated with propidium monoazide (PMA, Biotium, Fremont, CA, USA), a photoreactive dye that effectively eliminates PCR amplification of DNA from dead cells, thus providing discrimination of live versus dead bacteria by the viability PCR assay. PMA was prepared based on our previous study (19) in which 1 mg of PMA was dissolved in 1.9 ml of 20% DMSO (Sigma-Aldrich) to obtain a 1 mM stock solution. The stock preparation was stored in the dark at -20°C. 25 µL of PMA working stock solution was added to 500 µl of each previously prepared bacterial cell lysate to obtain a final concentration of 50 μM. Tubes were vortexed to ensure homogeneity of the solution, then incubated in the dark at room temperature for 10 minutes on a rocker. The microtubes were then placed horizontally in a PMA-Lite LED Photolysis Unit (Biotium, Fremont, CA) and exposed to LED light for 15 min, allowing for crosslinking of the PMA dye with bacterial DNA. Thereafter, the microtubes were centrifuged at 14,000 × g for 10 min. The supernatants were carefully discarded, and the cell pellets were stored at -20°C until used or processed immediately for DNA extraction (24).

#### 2.5.3. DNA purification

The cell pellets obtained after PMA treatment were lysed, and total genomic DNA extracted and purified as described in (15). DNA yield was then measured by a NanoDrop® ND-1000 Spectrophotometer (Thermo Fisher Scientific Inc., Waltham, USA) using A260/280 ratio with 1 μL of sample. Samples were then diluted to a final 1 ng/μL working dilution.

To generate the standard curve in the qPCR reaction, 4 ×10^7^ BCG (grown to log phase) were also lysed, and DNA extracted as above. DNA eluates were stored at -20oC until used or processed immediately for qPCR.

#### 2.5.4. TaqMan Quantitative Real-Time PCR

TaqMan quantitative Real-Time PCR targeting the lpqT (Rv1016c) gene was used to measure the viable BCG within the 24 hrs E/CTL co-cultures as described in (19), with some modifications. The lpqT gene belongs to a group of lipoprotein genes that are present in all bacteria (25) and is thought to be required for optimal growth of mycobacteria in vivo (26). In Brief, each reaction contained 2× master mix (12.5 µL), forward and reverse lpqT primers (0.5 µM final conc.), FAM labeled lpqT probe (0.2 µM final conc.), template DNA (5 µl = 5 ng) and nuclease-free dH_2_O to a final volume of 25 µl. The cycling parameters consisted of 10 min incubation at 95°C to activate the Taq, 40 cycles of 95°C for 10 s and 60°C for 30 s, followed by a cooling step at 40oC for 10 s. The temperature transition rate for all cycling steps was 20°C/s. The mean cycle threshold (CT) values of 3 replicates were analyzed using StepOne Software v2.1 (Applied Biosystems, CA) and used to calculate the percent of viable bacteria present in each sample. The PCR data were considered valid when the following criteria were achieved: PCR efficiency between 90 and 110%, regression coefficient (R2) value > 0.99, and the standard deviation ≤ 0.250. The final CT value in each treatment, considered for statistical analysis, was the average mean of two technical replicates for each of the two biological experiments. The final averages of all CT values for BCG and BCG*rel* cell cultures, from BCG and BCG*rel* vaccinated steers respectively, were compared to the final CT average obtained with the unvaccinated steers. CT values are inversely proportionate to the viable bacteria in the tested samples which rely upon a DNA-intercalating, PCR-inhibiting PMA treatment with qPCR output to define the viable bacteria.

### 2.6. Statistical analysis

A full factorial generalized linear mixed model analysis was conducted on each dataset using the SAS software procedure PROC GLIMMIX (SAS Institute Inc., Cary, NC, USA). Proportional responses (proliferative, cytokine-positive, and lethal signal-positive) were modeled using the beta distribution. The average CT value response was best fit as gaussian. The improved method of Kenward and Roger (DDFM=KR2) was used to compute the denominator degrees of freedom for the tests of fixed effects. Each model included a random intercept term for subjects and used a simple diagonal covariance structure (TYPE=VC).

Analyses of the proliferative and cytokine response data included fixed effect terms for vaccination group (BCG*rel* vaccinated, BCG vaccinated, unvaccinated), type of memory T cell (CD4 or CD8), post vaccination week (second or sixth), in vitro stimulus condition (BCG*rel*, BCG, or mock), and, for the latter dataset, the cytokine measured (IFN-γ, IL17A, IL-22, and TNF-α). Each model also included a random intercept term for the interaction of postvaccination week with steer nested within vaccination group. The analysis of the lethal signal response data included fixed effect terms for vaccination group (BCG*rel* or BCG vaccinated), type of memory T cell (CD4 or CD8), in vitro stimulus condition (BCG or BCG), and for the lethal signal measured, and a random intercept term for steer within *rel* vaccination group. The analysis of the average CT response included fixed effect terms for vaccination group (BCG*rel* vaccinated, BCG vaccinated, unvaccinated) and ex vivo stimulus condition (BCG*rel*, BCG, or mock), and a random intercept term for steer within vaccination group.

Significant model terms (P < 0.05) were analyzed post hoc by forming specific contrasts of interest and using the multiple comparisons technique of Holm (step-down Bonferroni; PHolm < 0.05). Modeled responses are summarized in figures as the predicted least squares means with 95% confidence intervals.

## 3. Results

### 3.1 Initial studies

Initial studies conducted ex vivo with cattle showed two rounds of stimulation of PBMC with APC primed with antigens were required to obtain enough cells for analysis (15). Currently, there is no way to examine the early events of Ag processing and presentation to CD4 and CD8 T cells in vivo. The first opportunity to examine the immune response elicited by a vaccine is through examining memory T cells that appear in blood after vaccination. A general method of vaccination of cattle is through an intradermal injection of a vaccine in the neck. Some of the vaccine is taken up by APC present in the tissue and transported to lymph nodes through the lymphatics where antigen processing and presentation takes place. Some of a vaccine may be transported by lymph to the regional lymph nodes draining the tissue where the vaccine is taken up by resident APC and processed. The time when a sufficient number of Ag specific memory T cells have proliferated and entered the blood stream, has not been determined. Two pilot studies were conducted to establish a time point when sufficient memory T cells were consistently present in blood to detect a recall response in all animals under study. Studies with 2 steers remaining from the ex vivo studies were vaccinated and tested to determine the earliest time point when a recall response could be consistently detected. The results indicated 14 days was the earliest time point to check. Two additional steers were used to verify this was a good time point for studying the recall response to vaccination (data not shown). Based on these observations a formal study was conducted with a larger number of steers. We were interested in the same parameters examined in the ex vivo studies. We were interested in determining if differences could be detected in the CD4 and CD8 memory T cell recall response in presence and absence of gene products encoded by genes under the regulation of *rel*.

### 3.2. Comparison of the proliferative recall responses of CD4 and CD8 memory T cells steers vaccinated with BCG*rel* or with BCG at 2 and 6 weeks PV

Three time points, 2, 4 and 6 weeks, were selected for determining if differences could be detected in the proliferative immune response between steers vaccinated with BCG*rel* in comparison with BCG over time. The data obtained at 4 weeks were essentially the same as results obtained at 6 weeks. Therefore, only the data obtained at 6 weeks are described. To increase the potential of detecting a difference in the proliferative response elicited by APC primed with BCG*rel*, or with BCG, PBMC from steers vaccinated with BCG*rel* were stimulated with APC primed with BCG*rel* or with BCG. PBMC from steers vaccinated with BCG were treated the same.

No significant differences were observed in the proliferative response of memory CD4 or CD8 T cells in PBMC from steers vaccinated with BCG*rel* stimulated with APC primed with BCG*rel* or BCG. Results with PBMC from steers vaccinated with BCG were the same at 2 and 6 weeks. However, the proportions of CD4 and CD8 memory T cells increased by 6 weeks. There was an equivalent increase in the CD4 and CD8 proliferative response (Figs. 2 and 3).

**Figure 2.**
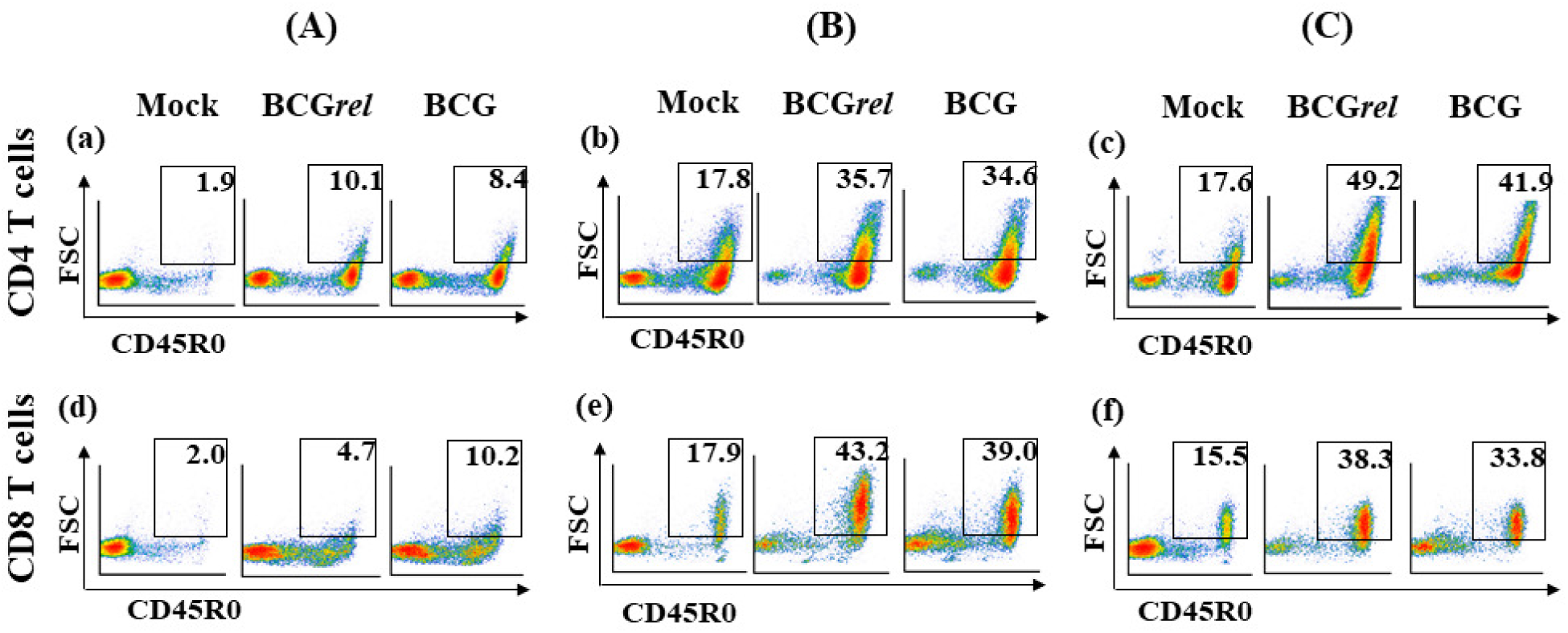
Representative flow cytometric (FC) profiles obtained with PBMC from a single steer showing how activated CD4 and CD8 memory T cells were visualized to obtain data for statistical analysis of the PBMC proliferative response: (A) FC profiles of ^a^CD4 and ^d^CD8 memory T cells from a control unvaccinated steer cultured alone or in the presence of either BCG*rel* or BCG. (B) FC profiles of ^b^CD4 and ^e^CD8 memory T cells from a steer vaccinated with BCG*rel* cultured alone or in the presence of either BCG*rel* or BCG. (C) Profiles of ^c^CD4 and ^f^CD8 memory T cells from a steer vaccinated with BCG cultured alone or in the presence of either BCG*rel* or BCG.

**Figure 3.**
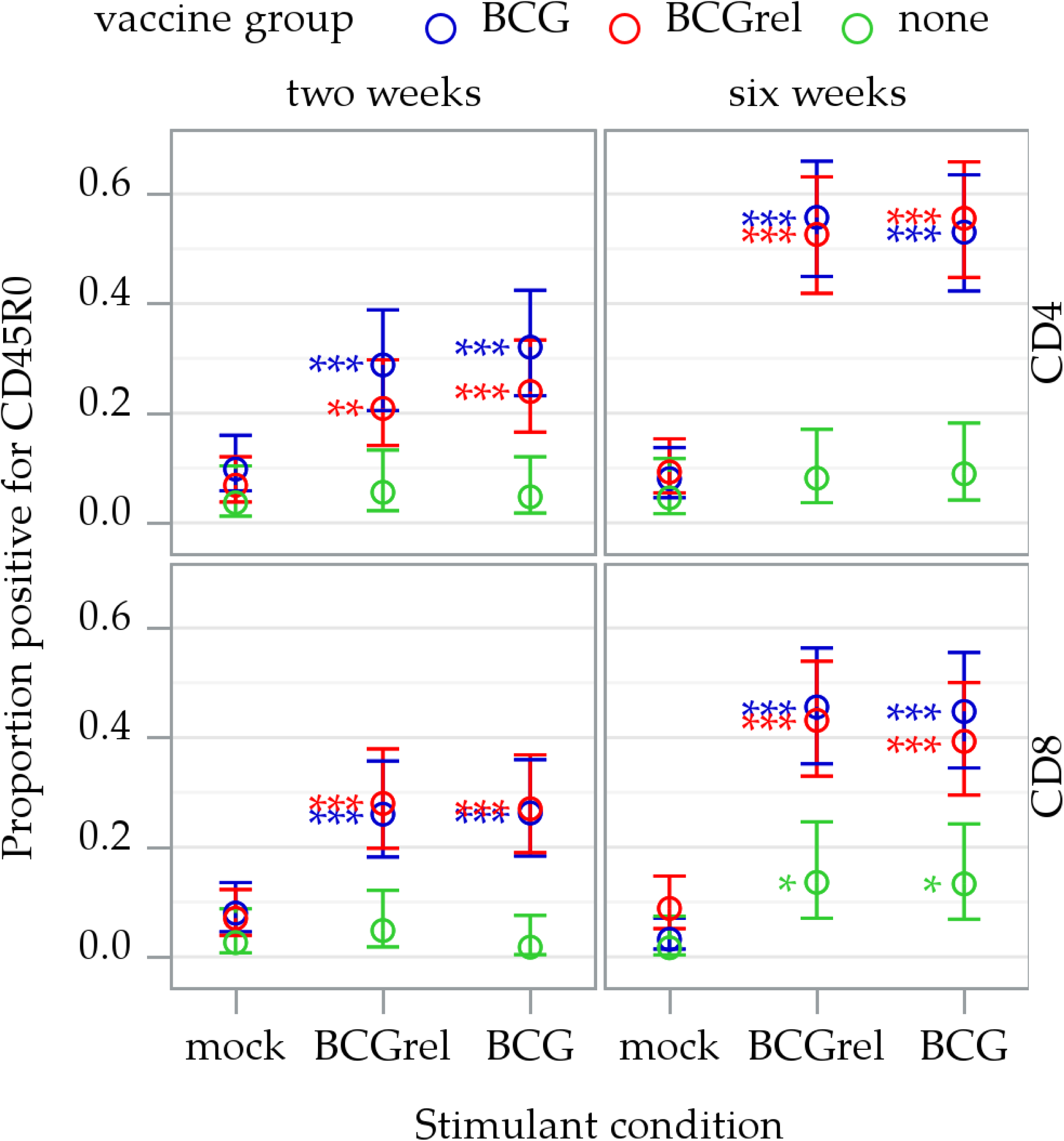
Summary of the proliferative recall responses of CD4 and CD8 memory T cells at 2, and 6 weeks PV, following stimulation ex vivo. The data were obtained from 2 replicates for each time point. The PBMC were collected at post vaccination weeks 2, 4, and six from steers vaccinated with BCG (n = 4), BCG*rel* (n = 4), or left unvaccinated (n = 2). Since the data obtained at 4 weeks was essentially the same as 6 weeks, only the data from weeks 2 and 6 are presented. The data shown are the predicted means and 95% confidence intervals resulting from a full factorial generalized linear mixed model analysis. Stimulant-induced change is the difference between ex vivo stimulation (with BCG*rel* or BCG) and “mock” stimulation. Significance within vaccine groups is indicated by color-matched asterisks (*, PHolm < 0.05; **, PHolm < 0.001; ***, PHolm < 0.0001). No significant differences were observed at 2 or 6 weeks PV.

### 3.3. Comparison of expression of IFN-γ, TNF-α, IL-17A, and IL-22 in CD4 and CD8 memory T cells in PBMC from steers vaccinated with BCG*rel* or BCG

Studies with IFN-γ, TNF-α, and IL-17A in our initial ex vivo studies showed there was no detectable differential in expression in CD4 and CD8 T cells associated with stimulation with APC primed with BCG*rel* or with BCG. They were all upregulated. Ex vivo, there was limited opportunity for BCG to express any gene products before being processed for Ag presentation. The study was repeated looking at development of an immune response in vivo where there was time for expression of bacterial products that might modulate development of CD4 and CD8 memory T cells. Studies were conducted to determine if any differences could be detected in the memory CD4 and CD8 T cells proliferating in response to stimulation with either BCG*rel* or BCG. IL-22 (another cytokine implicated in regulating the immune response to mycobacteria was included in the study (cited in (18, 27)). To increase the potential of detecting a difference in expression of the cytokines in PBMC from steers vaccinated with BCG*rel,* PBMC were stimulated with APC primed with BCG*rel* or with BCG. PBMC from steers vaccinated with BCG were treated the same. Comparison of expression of IFN-γ with expression of TNF-α, IL-17, and IL-22 revealed no significant difference in the proportion of CD4 and CD8 T cells expressing the cytokines. All the cytokines were coexpressed in the same cell subsets of CD4 and CD8 memory T cells. Since no significant differences in expression of the cytokines were observed in CD4 or CD8 T cell subsets, the data were combined for presentation in a single figure (Figs. 4 and 5).

**Figure 4.**
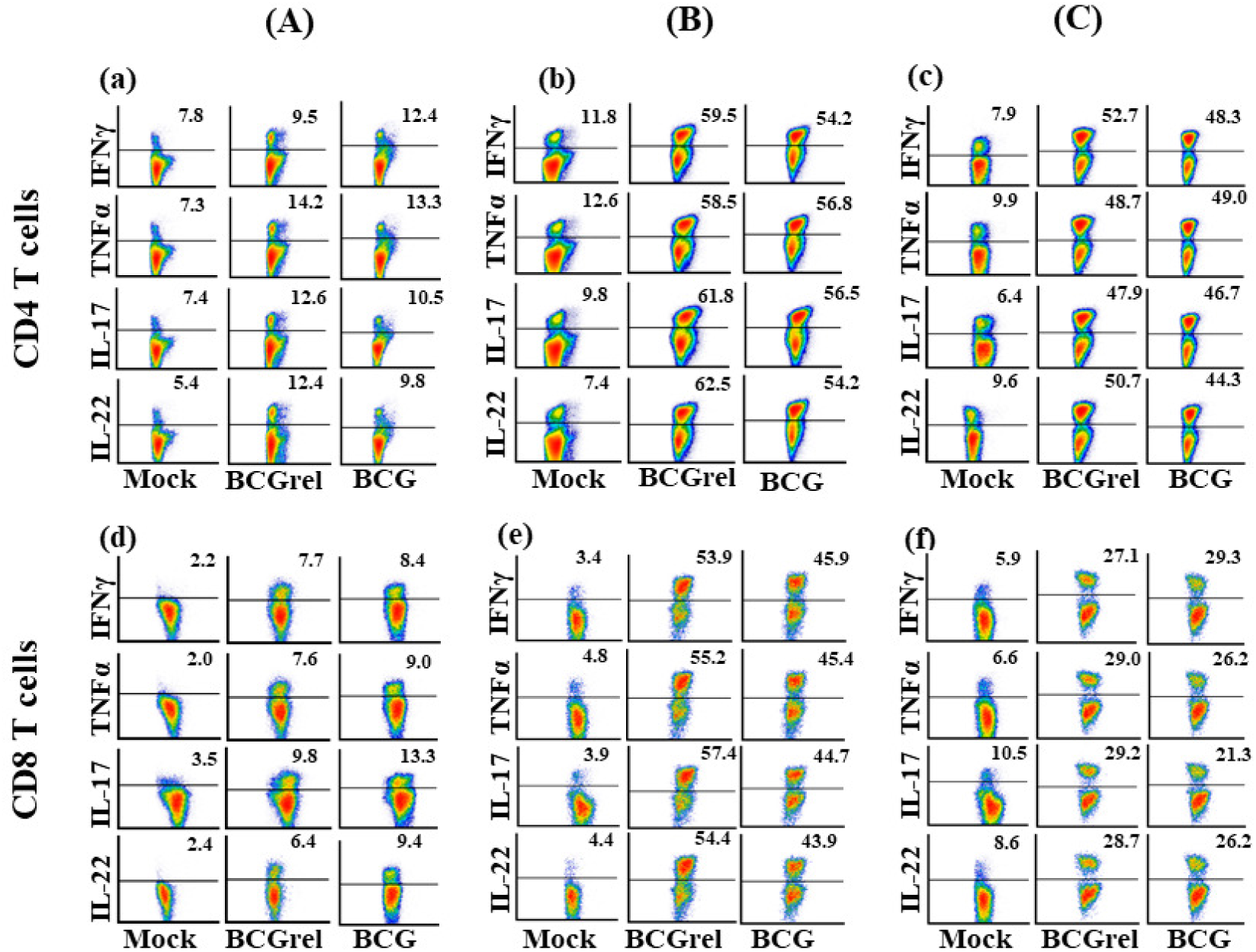
Representative flow cytometric (FC) profiles obtained with PBMC from a single steer showing how activated CD4 and CD8 memory T cells were visualized to obtain data for statistical analysis of expression of IFN-γ, TNF-α, IL-17, IL-22. (A) FC profiles of ^a^CD4 and ^d^CD8 memory T cells from a control unvaccinated steer cultured alone or in the presence of either BCG*rel* or BCG. (B) FC profiles of ^b^CD4 and ^e^CD8 memory T cells from a steer vaccinated BCG*rel* cultured alone or in the presence of either BCG*rel* or BCG. (C) Profiles of ^c^CD4 and ^f^CD8 memory T cells from a steer vaccinated with BCG cultured alone or in the presence of either BCG*rel* or BCG.

**Figure 5.**
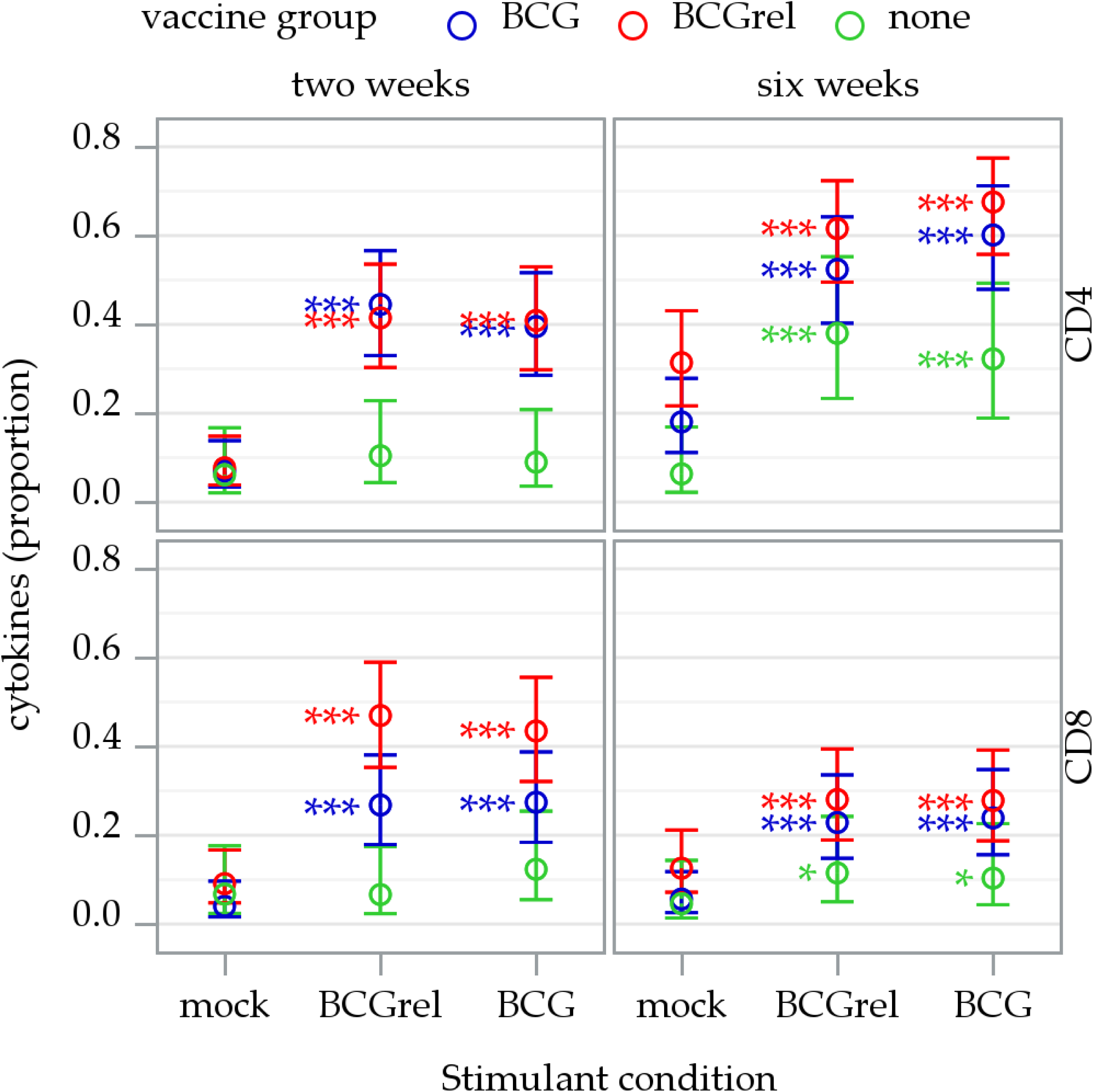
Summary of cytokine expression by CD4 and CD8 memory T cells at 2 and 6 weeks PV. The data were obtained from 2 replicates for each time point from control unvaccinated steers (n = 2) and steers vaccinated with BCG (n = 4) or BCG*rel* (n =4) showing the proportion of activated CD4 and CD8 memory T cells expressing IFN-γ, TNF-α, IL-17A, and IL-22 in PBMC cultures left unstimulated (mock) or stimulated with BCG or BCG*rel*. The results obtained from the individual comparisons were essentially the same. So, the data were combined so that all the relevant data could be presented in a single graph with an example of the data from an individual steer shown in Fig. 4. The data shown are the predicted means and 95% confidence intervals resulting from a full factorial generalized linear mixed model analysis. Stimulant induced change is the difference between ex vivo stimulation (with BCG*rel* or BCG) and “mock” stimulation. Significance within vaccine groups is indicated by color-matched asterisks (*, PHolm < 0.05; **, PHolm < 0.001; ***, PHolm < 0.0001).

### 3.4. Comparison of the memory CD8 CTL response that develops in steers vaccinated with BCG*rel* or BCG

Multiple studies with candidate proteins and deletion mutants in *Mtb, Mbv*, and BCG have shown the immune responses, that develop, modulate the immune response in mouse models and cattle. However, none have elicited an immune response that prevents establishment of a persistent infection. Mutants with a deletion in *rel* have shown ability to establish a persistent infection is associated with the presence of gene products encoded by genes under the regulation of *rel*. How and when these products interfere with immune clearance has not been determined. Analysis of the primary immune response ex vivo demonstrated that a comparable CD8 CTL response develops following stimulation with APC primed with BCG or with BCG*rel*. As mentioned, there was limited opportunity for BCG or BCG*rel* to synthesize any gene products ex vivo before being processed by APC for Ag presentation to CD4 and CD8 T cells. Comparison of the memory CD4 and CD8 CTL recall response with PBMC from steers vaccinated with either BCG*rel* or BCG revealed the gene products do not interfere with the timing of appearance of an immune response as detected by proliferation of CD4 and CD8 memory T cells at 14 days PV (Figs. 2 and 3). Comparison of the proportion of CD4 and CD8 memory T cells that express 4 different cytokines involved in regulation of the immune response did not reveal any detectable differences in expression of molecules involved in regulation of the immune response. As part of the studies the same steers were used to determine if any difference in functional activity could be detected in the CD4 or CD8 memory T cells that develop following vaccination with BCG*rel* or with BCG. PBMC from steers vaccinated with either BCG*rel* or BCG were stimulated with APC primed with BCG*rel* or with BCG. The whole study was conducted 2 times 2 days apart at 6 weeks. Blood was collected from all the steers at 6 weeks and 2 days later.

Comparison of the memory CD8 CTL recall response with PBMC from the steers vaccinated with BCG*rel* and stimulated with APC primed with BCG*rel* or with BCG showed stimulation with BCG*rel* elicited a much more vigorous CD8 CTL response than stimulation with BCG (Figs. 6 and 7). The response to stimulation with APC primed with BCG was not significant. Comparison of the memory CD8 CTL response with PBMC from steers vaccinated with BCG stimulated with BCG*rel* showed the CD8 CTL response was significant with APC primed with BCG*rel* but less than the response observed with CD8 CTL from the steers vaccinated with BCG*rel*. The CD8 CTL response of PBMC from steers vaccinated with BCG with stimulation withto APC primed with BCG was not significant (Figs. 6 and 7)

**Figure 6.**
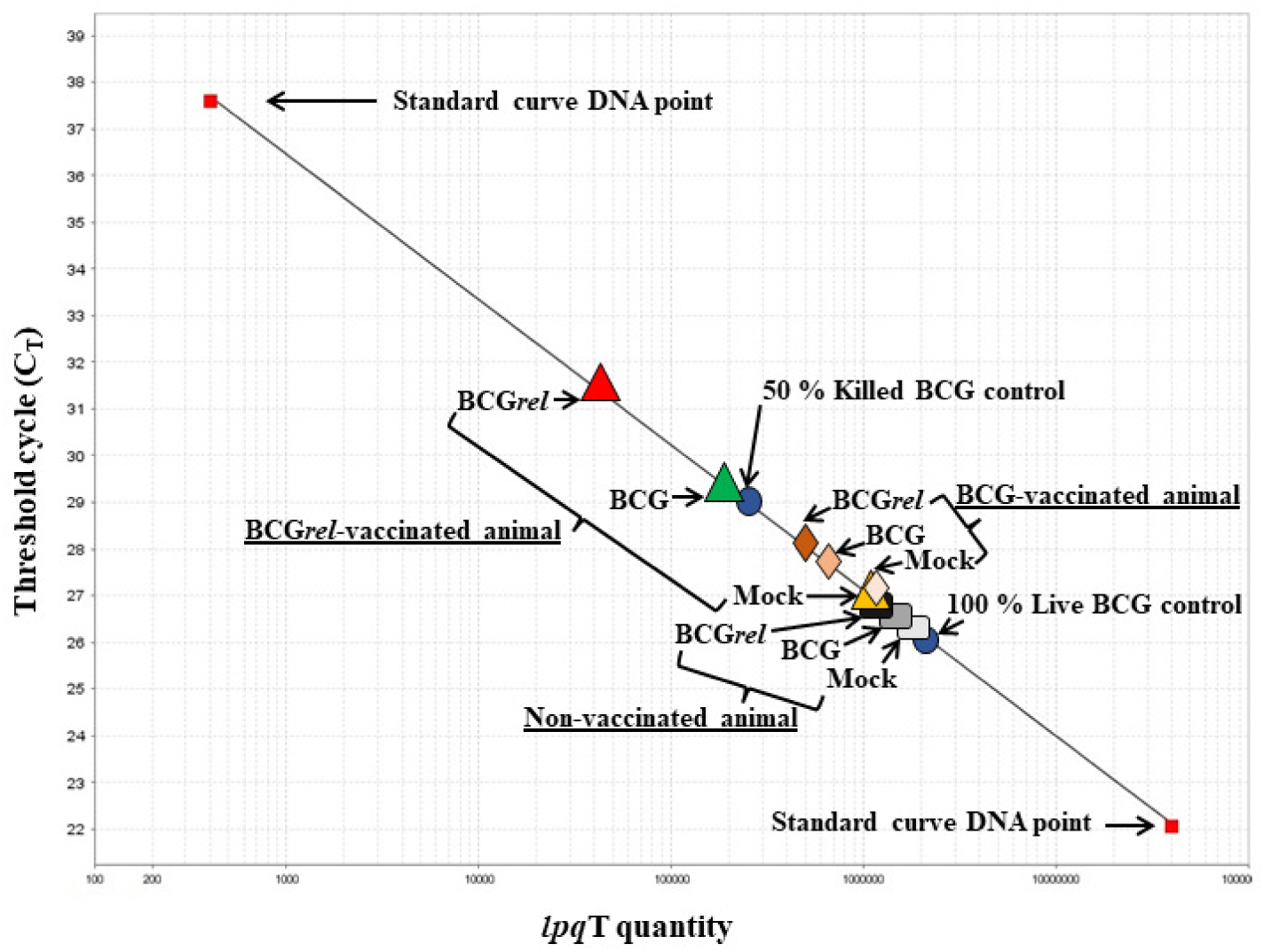
Representative standard DNA curve developed and used to obtain data on intracellular killing of BCG in MoMac target cells from one animal of each vaccination group. Standard curve with 8 dilutions of DNA from pure live BCG (red squares) 4 × 10^7^ to 4 copies. <u>Blue oval data points</u> represent two DNA viability controls prepared from 100% live (the lower blue oval) and from mixtures of 50% live/50% dead bacteria (the upper blue oval) after PMA treatment. <u>CT values</u> represent averages of triplicate preparations of DNA from infected target MoMac co-cultured with BCG*rel* or BCG stimulated T cells or left unstimulated (mock) from BCG*rel* vaccinated animal are <u>represented by red, green, and yellow triangles respectively</u>. <u>The culture matched CT values</u> from BCG vaccinated animal are <u>represented by dark, light, and faint light orange diamonds respectively</u>. <u>The culture matched CT values</u> from a control unvaccinated animal <u>are represented by dark, light, and faint light blue rectangles respectively</u>.

**Figure 7.**
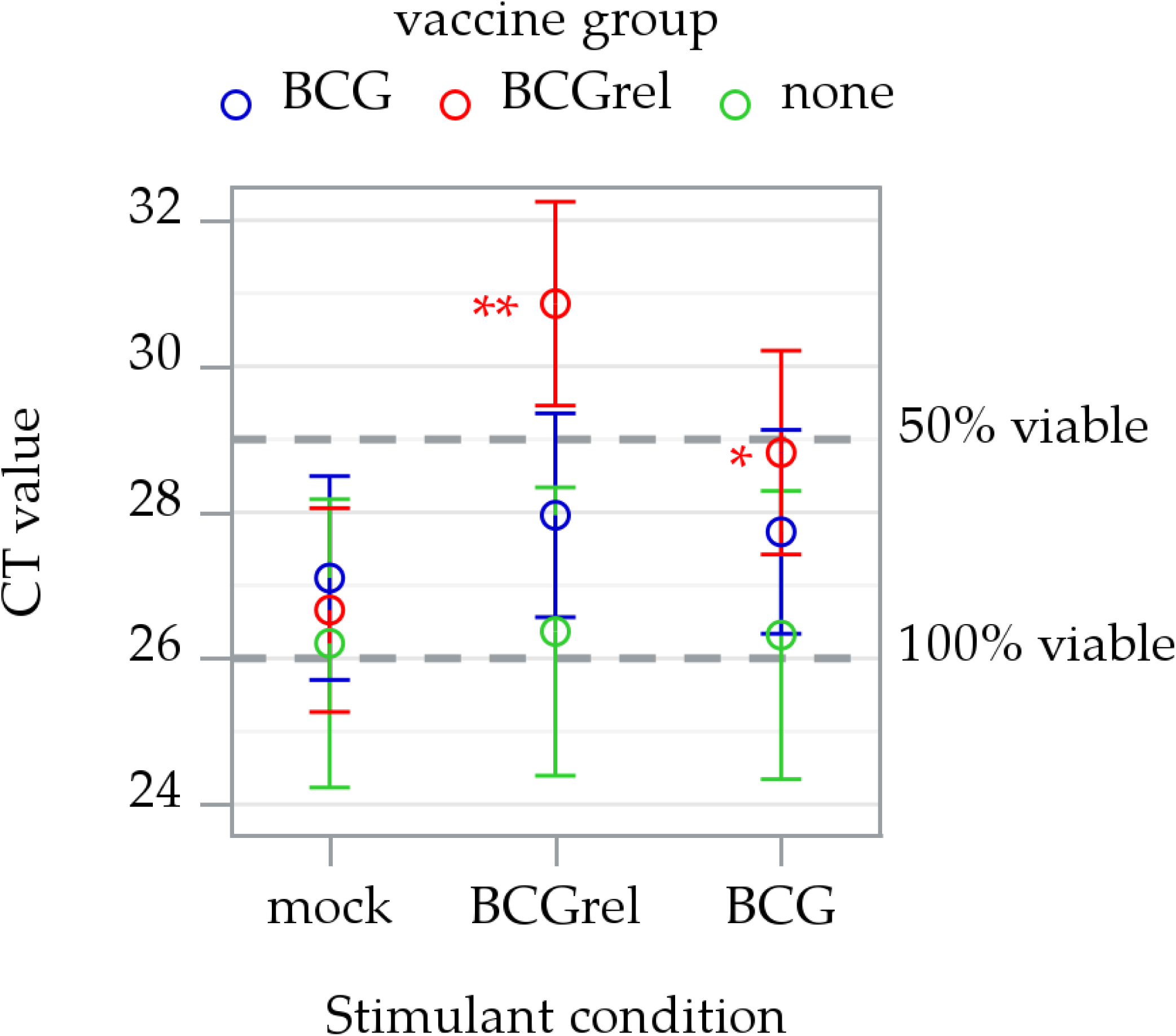
Summary of bacterial killing mediated by CD8 CTL 6 weeks PV, using PMA/qRTPCR. The data were obtained from 2 replicates of control unvaccinated steers (n = 2) and steers vaccinated with BCG (n = 4) or BCG*rel* (n = 4) showing the extent of intracellular killing of bacteria in infected target cells overlaid with cultured PBMC left unstimulated (mock) or stimulated with BCG*rel* or BCG. Intracellular bacterial viability was measured at 24 hours, where CT values of ∼26 or less correspond to 100% viable bacteria and a value ∼29 corresponds to 50% viable bacteria. The data shown are the predicted means and 95% confidence intervals resulting from a full factorial generalized linear mixed model analysis. Stimulant-induced change is the difference between ex vivo stimulation (with BCG*rel* or BCG) and “mock” stimulation. Significance within vaccination groups is indicated by color-matched asterisks (*, PHolm < 0.05; **, PHolm < 0.001; ***, PHolm < 0.0001).

### 3.5. Comparison of the content of perforin, GranzB, and granulysin in memory CD4 and CD8 CTL proliferating in response to stimulation with APC primed with BCG*rel* or with BCG

The observation that the functional activity of memory CD8 CTL was less in cells stimulated with APC primed with BCG than with APC primed with BCG*rel* suggested the difference might be associated with a difference in the content of one or all the molecules involved in intracellular killing of bacteria. As a follow up of the killing assay, blood was collected at 8 weeks PV to determine if there was any detectable difference in the content of perforin, GranzB, or granulysin. The proportion of cells containing perforin, GranzB, and granulysin was used as an indirect measure of the content of each molecule. The same labeling format was used for labeling to identify any difference in the immune response elicited by vaccination with BCG*rel* or with BCG. No significant difference was observed in the proportion of perforin, GranzB, or granulysin in CD4 or CD8 T cells (Figs. 8 and 9).

**Figure 8.**
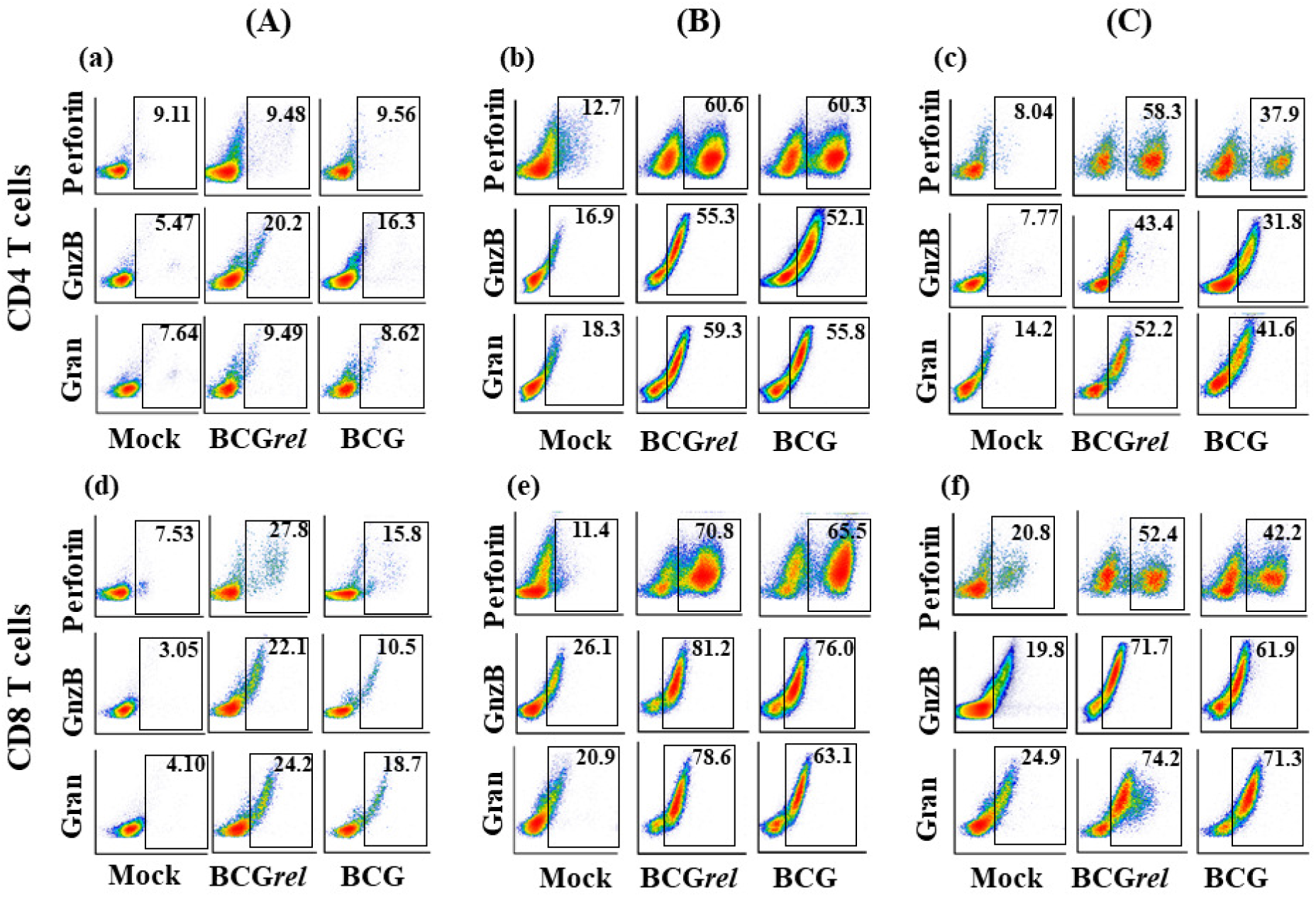
Representative flow cytometric (FC) profiles obtained with PBMC from a single steer showing how activated CD4 and CD8 memory T cells were visualized to obtain data for statistical analysis of expression of perforin, GranzB, and granulysin in CD4 and CD8 memory T cells. (A) FC profiles of ^a^CD4 and ^d^CD8 memory T cells from a control unvaccinated steer cultured alone or in the presence of either BCG*rel* or BCG. (B) FC profiles of ^b^CD4 and ^e^CD8 memory T cells from a steer vaccinated BCG*rel* cultured alone or in the presence of either BCCG*rel* or BCG. (C) Profiles of ^c^CD4 and ^f^CD8 memory T cells from a steer vaccinated with BCG cultured alone or in the presence of either BCG*rel* or BCG.

**Figure 9.**
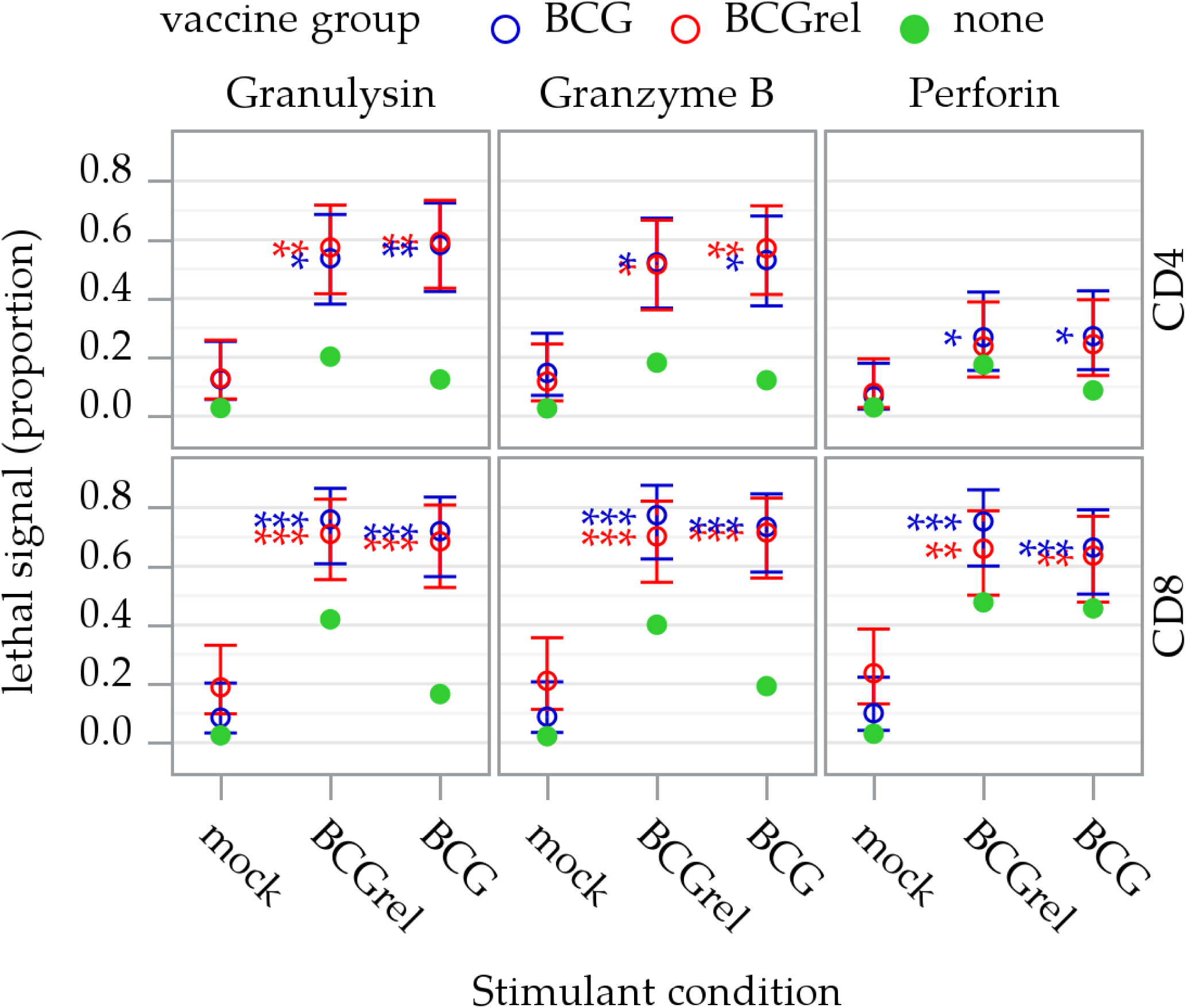
Summary of data obtained at eight weeks from 3 replicates of a control unvaccinated steer (n = 1) and steers vaccinated with BCG (n = 2) or BCG*rel* (n = 2) showing the proportion of perforin, GranzB, and granulysin containing CD4 and CD8 memory T cells in PBMC cultures left unstimulated (mock) or stimulated with BCG or BCG*rel*. The data shown for vaccinated steers (open circles) are the predicted means and 95% confidence intervals resulting from a full factorial generalized linear mixed model analysis. The data for the single unvaccinated steer are shown as closed circles. Stimulant-induced change is the difference between ex vivo stimulation (with BCG*rel* or BCG) and “mock” stimulation. Significance within vaccinated steer groups is indicated by color-matched asterisks (*, PHolm < 0.05; **, PHolm < 0.001; ***, PHolm < 0.0001). No significant difference was observed in the proportion of memory CD4 or CD8 T cells containing the molecules that transmit the lethal signals associated with intracellular killing.

## 4. Discussion

Extensive studies have been conducted with mice (8, 28–33), ruminants (34–38), non-human primates (39), and humans (40) during the past 30 plus years to characterize the immune response to mycobacterial pathogens and develop vaccines. The major impediment encountered in development of vaccines against pathogenic mycobacteria has been lack of understanding of the mechanisms used by the pathogens to evade clearance by the immune system and establish a persistent infection. Studies, primarily with *Mtb, Mbv*, and *Map*, have shown the immune response can be modulated by vaccination, but none of the methods devised to develop vaccines have yielded a vaccine that prevents establishment of a persistent infection (8–10). These efforts have included attempts to improve the efficacy of BCG as a vaccine by deleting additional genes associated with virulence.

Studies of the stringent response in mice have provided the first lead to identifying the mechanisms used by mycobacterial pathogens to evade immune clearance. In the absence of *rel*, the ability of mycobacteria to establish a persistent infection is lost (7). The observation indicated that genes under the regulation of *rel* include genes encoding products that interfere with immune clearance. The development of assays to study the primary and recall responses in real time, using the same tissue culture platform, provided an opportunity to begin studies focused on determining how gene products encoded by genes regulated by *rel* interfere with immune clearance of mycobacterial pathogens. The methods can be used with any species including humans to replicate the studies described in our initial and present report (41).

The first observation of global importance gained from the study of *rel* deletion mutants is that deletion has the same effect on the 3 mycobacterial pathogens affecting humans and animals *Mtb*, *Mbv*/BCG, Map. It can be predicted that deletion will have the same effect on other pathogenic mycobacteria. The additional observations provide information on the immune response to mycobacterial pathogens. Studies of the primary immune response ex vivo show the immune response is MHC restricted and that concurrent Ag presentation by MHC I and MHC II molecules to CD4 and CD8 T cells is essential for development of CD8 CTL for mycobacterial pathogens (19, 42). These observations are consistent with previous studies (reviewed in (43)). The added information is the demonstration that the development of CD8 CTL requires the concurrent interaction of CD4 and CD8 T cells with APC at the time of Ag presentation. Comparison of the CD4 and CD8 T cell primary immune responses that develop following stimulation with APC primed with BCG or with BCG*rel* ex vivo in tissue culture show there is no detectable difference in the development of CD8 CTL able to kill intracellular bacteria. Likewise, there are no differences in expression of IFN-γ, TNF-α, or IL-17 cytokines known to play a role in regulating the immune response. Of interest and contrary to some comparative studies of expression of these cytokines in other species, there is no difference in expression of these cytokines in responding CD4 and CD8 T cells. The results were the same for IL-22. A mAb specific for this cytokine was not available at the time or our initial studies. The current study was not designed for further in depth studies of these cytokines. The study was designed to try and reveal any difference in the immune response to BCG*rel* and BCG. A difference in the functional activity was detected. Further studies revealed the reduction in functional activity was not associated with content of proteins involved in intracellular killing of bacteria, perforin, GranzB, and granulysin.

In the ex vivo studies, there was limited time for BCG to synthesize products involved in regulating the immune response. Under these experimental conditions antigen processing and presentation of BCG and BCG*rel* by APC leads to development of CD8 CTL with equal ability to kill intracellular bacteria. The data indicate no gene products are present in cultured BCG that interfere with Ag processing and presentation by APC.

There has been no opportunity to look at the primary immune response in vivo. Thoughts on what is believed to be happening in the in vivo environment have been derived from studies with mycobacterial pathogens with an intact *rel* gene. The results obtained in the studies have been informative and provide background for interpretation of the results obtained in the present studies. Results from study of events occurring in the lung after aerosol exposure have shown there is mixed signaling occurring during establishment of a persistent infection and development of granulomas (31, 44). Bacteria are taken up by multiple cell types including dendritic cells, alveolar, interstitial macrophages, and neutrophils (45). Positive and negative signaling may be derived from different cell types with some signaling associated with initiation of granuloma formation. Studies by Grant, Flynn et al. in non-human primates using PET CT imaging have obtained information showing the fate of tagged bacteria introduced into the lung (46). This has provided a visual record of the events leading to development of granulomas. The transient appearance of some granulomas and persistence of others suggest the involvement of innate and adaptive immune responses. Products produced by mycobacteria could be playing a role in interfering with immune clearance, allowing for persistence of some bacteria and development of some granulomas and clearance of other granulomas. Although indirect, data from a recent comprehensive study of granulomas in humans at different stage of disease have provided additional information on the organization and spatial distribution of cell types in granulomas (47). Of interest, 2 molecules associated with regulation of immune function were identified that could be playing a role in modulating the function of CD8 T cells, PD-L1 and IDO1 (48, 49). None of the studies have provided opportunity to directly study the function of CD4 and CD8 T cells that develop in the presence of the gene products produced by mycobacteria in vivo. The studies by Dahl et al. have shown the initial events of infection and granuloma formation still occur but resolve over time, suggesting an immune response needs to develop. The clearance of the granulomas is associated with clearance of bacteria. Granuloma formation was not examined in the present study. Studies in cattle have shown granulomas do develop in lungs following experimental infection (50, 51). It was inferred from the study in mice and results from the study with *Map* that the BCG*rel* deletion mutant was cleared. Exactly what is occurring during the in vivo environment in the lung and other tissues will be difficult to sort out. Follow up studies of the memory recall response, as described in the present report, have provided the opportunity to analyze the development of the memory recall response in the presence and absence of gene products produced by genes under the regulation of *rel*. The memory T cells that appear in the blood provide a summation of events that have occurred after initial exposure. The comparison of the proliferative response revealed gene products under the regulation of *rel* do not interfere with initiation of an immune response as reflected by the timing of a detectable proliferative response. Comparison of cytokines involved in regulation of the immune response revealed no difference in the expression of IFN-γ, TNF-α, IL-17, or IL-22. Comparison of the content of molecules involved in the killing of intracellular bacteria revealed no difference in memory T cells from steers vaccinated with BCG*rel* or with BCG. Comparison of the functional activity revealed a difference. This was revealed by cross comparing the memory CD8 CTL responses elicited by APC primed with BCG*rel* or with BCG. Comparison of the recall response with PBMC from steers vaccinated with BCG*rel,* using APC primed with BCG*rel* or with BCG, showed stimulation with BCG*rel* elicited a vigorous CTL response. In contrast stimulation with BCG elicited a response that was not significant. Comparison of the recall response from steers vaccinated with BCG showed stimulation with APC primed with BCG*rel* elicited a CD8 CTL response that was significant but less than the response observed with PBMC from steers vaccinated with BCG*rel*. Stimulation with APC primed with BCG elicited a response that was not significant. The study of the memory recall response revealed development of functional activity was impaired in vivo in the presence of gene products produced when *rel* is present.

To our knowledge, we are the first investigators to recognize that genes encoding products regulating the immune response are included in the genes regulated by *rel*. Research on the stringent response, conducted thus far, has focused on characterization of the global effect of interfering with synthesis of (p)ppGpp, the alarmone synthesized or hydrolyzed by Rel and Rel homologues. Control of expression of an array of gene products is lost by disrupting *rel* homologues in different lineages of bacteria (reviewed in Atkinson et al. (52)). Information on the role of (p)ppGpp in virulence of Gram-negative and Gram-positive bacteria and plant pathogens is reviewed in Dalebroux (53). A more recent review by Kundra et al. extends information in the review by Dalebroux et al. and provides further information on the role of (p)ppGpp in regulating the stringent response in the vertebrate host (54). The review summarizes data showing the pivotal role of (p)ppGpp in regulating expression of gene products associated with virulence, antibiotic resistance, and persistence in different lineages of bacteria. Limited but important studies of other pathogens have shown deletion of the homologues of *rel* in other pathogenic bacteria has the same effect: Brucella sp (55), Francisella (56), Borrelia (57) Listeria (58) S. enterica (59), S. typhimurium (60), and Staphylococcus (61). The more comprehensive studies with mycobacterial pathogens have provided data that show some of the genes regulated by *rel* synthesize products that interfere with the function of the immune system. Further studies are clearly needed to duplicate and extend the information obtained from the present study. The replacement of the chromium release assay and the CFU assay have provided opportunity to study the immune response to mycobacterial pathogens in real time. The methods developed to study the primary and recall responses ex vivo can be used with any species including humans (41). The use of cattle in the present studies has highlighted the value of including a large animal species in the study of mycobacterial pathogens (18, 62).

## 5. Summary and conclusions

The results obtained in our initial and follow up investigations provide data showing adaptation of mycobacterial pathogens for survival in the vertebrate host include development of mechanisms to interfere with immune clearance. Comparative studies of the immune response and development of CD8 cytotoxic T cells against mycobacterial pathogens have revealed products encoded by genes regulated by *rel*, regulator of the stringent response, are involved. Studies with BCG in cattle show that, when *rel* is present, APC primed with BCG elicit development of CD8 CTLs able to control but not clear infection, leading to a persistent infection. When *rel* is deleted from the genome, APC primed with a *rel* deletion mutant elicit development of CD8 CTL able clear the infection. Identification of the gene products that interfere with immune clearance should lead to development of a vaccine that will elicit sterile immunity.

## Author Contributions

AHM, GSA, LMF, and WCD conceived the study. KTP developed the *rel* deletion mutants used to conduct the studies. AHM, GSA, KTP, and WCD participated in the design of the protocol to conduct the studies. AHM and GSA conducted the studies. VH, SEA participated in the conduct of the studies. AHM, GSA and DAS participated in statistical analysis of the data. AHM, GSA, WCD, KTP, LMF, WM, LB, NS, CDS and DAS participated in the writing and interpretation of the results. WCD obtained the funding for the project. WCD oversaw and participated in all aspects of the study. All authors read and approved the final manuscript.

## Acknowledgments

The authors wish to acknowledge the excellent technical support and animal care provided by Emma Karol and her staff. The senior author would like to recognize Sidney Raffel, Stanford University School of Medicine, mentor, and early pioneer studying mechanisms of pathogenesis of tuberculosis (1911–2013).

## References

1. Koch R. 1884. Die Aetiologie der tuberkulose. Mittheilungen ausdem Kaiserlichen Gesundheitsamte 2:1–88.

2. Johne HA, Frothingham L. 1895. Ein eigenthumlicher fall von tuberculosis beim rind [A peculiar case of tuberculosis in a cow]. Deutsche Zeitschr Tierm Path 21, 438–454:438–454.

3. Bachmann NL, Salamzade R, Manson AL, Whittington R, Sintchenko V, Earl AM, Marais BJ. 2019. Key Transitions in the Evolution of Rapid and Slow Growing Mycobacteria Identified by Comparative Genomics. Front Microbiol 10:3019.

4. Chung J, Ince D, Ford BA, Wanat KA. 2018. Cutaneous Infections Due to Nontuberculosis Mycobacterium: Recognition and Management. Am J Clin Dermatol 19:867–878.

5. Ratnatunga CN, Lutzky VP, Kupz A, Doolan DL, Reid DW, Field M, Bell SC, Thomson RM, Miles JJ. 2020. The Rise of Non-Tuberculosis Mycobacterial Lung Disease. Front Immunol 11:303.

6. Prasanna AN, Mehra S. 2013. Comparative phylogenomics of pathogenic and non-pathogenic mycobacterium. PLoS One 8:e71248.

7. Dahl JL, Kraus CN, Boshoff HIM, Doan B, Foley K, Avarbock D, Kaplan G, Mizrahi V, Rubin H, Barry CEI. 2003. The role of RelMtb-mediated adaptation to stationary phase in long-term persistence of Mycobacterium tuberculosis in mice. Proc Natl Acad Sci USA 100:10026–10031.

8. Dockrell HM, Smith SG. 2017. What Have We Learnt about BCG Vaccination in the Last 20 Years? Front Immunol 8:1134.

9. Lewis KN, Liao R, Guinn KM, Hickey MJ, Smith S, Behr MA, Sherman DR. 2003. Deletion of RD1 from Mycobacterium tuberculosis mimics bacille Calmette-Guerin attenuation. J Infect Dis 187:117–23.

10. Bannantine JP, Talaat AM. 2015. Controlling Johne’s disease: vaccination is the way forward. Front Cell Infect Microbiol 5:2.

11. Park KT, Dahl JL, Bannantine JP, Barletta RG, Ahn J, Allen AJ, Hamilton MJ, Davis WC. 2008. Demonstration of allelic exchange in the slow-growing bacterium Mycobacterium avium subsp. paratuberculosis, and generation of mutants with deletions at the pknG, relA, and lsr2 loci. Appl Environ Microbiol 74:1687–1695.

12. Walburger A, Koul A, Ferrari G, Nguyen L, Prescianotto-Baschong C, Huygen K, Klebl B, Thompson C, Bacher G, Pieters J. 2004. Protein kinase G from pathogenic mycobacteria promotes survival within macrophages. Science 304:1800–1804.

13. Park KT, Allen AJ, Bannantine JP, Seo KS, Hamilton MJ, Abdellrazeq GS, Rihan HM, Grimm A, Davis WC. 2011. Evaluation of two mutants of Mycobacterium avium subsp. paratuberculosis as candidates for a live attenuated vaccine for Johne’s disease. Vaccine 29:4709–19.

14. Park KT, ElNaggar MM, Abdellrazeq GS, Bannantine JP, Mack V, Fry LM, Davis WC. 2016. Phenotype and Function of CD209+ Bovine Blood Dendritic Cells, Monocyte-Derived-Dendritic Cells and Monocyte-Derived Macrophages. PLoS One 11:e0165247.

15. Abdellrazeq GS, Elnaggar MM, Bannantine JP, Park KT, Souza CD, Backer B, Hulubei V, Fry LM, Khaliel SA, Torky HA, Schneider DA, Davis WC. 2018. A Mycobacterium avium subsp. paratuberculosis relA deletion mutant and a 35 kDa major membrane protein elicit development of cytotoxic T lymphocytes with ability to kill intracellular bacteria. Vet Res 49:53.

16. Lurie MB. 1934. The Fate of Bcg and Associated Changes in the Organs of Rabbits. J Exp Med 60:163–78.

17. Palmer MV, Kanipe C, Cox R, Robbe-Austerman S, Thacker TC. 2019. Characteristics of subclinical Mycobacterium avium ssp. paratuberculosis infection in a captive white-tailed deer herd. J Vet Diagn Invest doi:10.1177/1040638719873028:1040638719873028.

18. Waters WR, Palmer MV, Thacker TC, Davis WC, Sreevatsan S, Coussens P, Meade KG, Hope JC, Estes DM. 2011. Tuberculosis immunity: opportunities from studies with cattle. Clin Dev Immunol 2011:11 pages.

19. Abdellrazeq GS, Mahmoud AH, Park KT, Fry LM, Elnaggar MM, Schneider DA, Hulubei V, Davis WC. 2020. relA is Achilles’ heel for mycobacterial pathogens as demonstrated with deletion mutants in Mycobacterium avium subsp. paratuberculosis and mycobacterium bovis bacillus Calmette-Guerin (BCG). Tuberculosis (Edinb) 120:101904.

20. Bardarov S, Bardarov SJ, Pavelka MSJ, Sambandamurthy V, Larsen M, Tufariello J, Chan J, Hatfull G, Jacobs WRJ. 2002. Specialized transduction: an efficient method for generating marked and unmarked targeted gene disruptions in Mycobacterium tuberculosis, M. bovis BCG and M. smegmatis. Microbiology 148:3007–3017.

21. Elnaggar MM, El-Naggar MM, Abdellrazeq GS, Sester M, Khaliel SA, Singh M, Torky HA, Davis WC. 2015. Development of an improved ESAT-6 and CFP-10 peptide-based cytokine flow cytometric assay for bovine tuberculosis. Comp Immunol Microbiol Infect Dis 42:1–7.

22. Elnaggar MM, Abdellrazeq GS, Dassanayake RP, Fry LM, Hulubei V, Davis WC. 2018. Characterization of alphabeta and gammadelta T cell subsets expressing IL-17A in ruminants and swine. Dev Comp Immunol 85:115–124.

23. Abdellrazeq GS, Elnaggar MM, Bannantine JP, Schneider DA, Souza CD, Hwang J, Mahmoud AHA, Hulubei V, Fry LM, Park KT, Davis WC. 2019. A peptide-based vaccine for Mycobacterium avium subspecies paratuberculosis. Vaccine 37:2783–2790.

24. Park KT, Allen AJ, Davis WC. 2014. Development of a novel DNA extraction method for identification and quantification of Mycobacterium avium subsp. paratuberculosis from tissue samples by real-time PCR. J Microbiol Methods 99:58–65.

25. Rezwan M, Grau T, Tschumi A, Sander P. 2007. Lipoprotein synthesis in mycobacteria. Microbiology 153:652–8.

26. Sassetti CM, Rubin EJ. 2003. Genetic requirements for mycobacterial survival during infection. Proc Natl Acad Sci U S A 100:12989–94.

27. Cowan J, Pandey S, Filion LG, Angel JB, Kumar A, Cameron DW. 2012. Comparison of interferon-gamma-, interleukin (IL)-17- and IL-22-expressing CD4 T cells, IL-22-expressing granulocytes and proinflammatory cytokines during latent and active tuberculosis infection. Clin Exp Immunol 167:317–29.

28. Perez E, Samper S, Bordas Y, Guilhot C, Gicquel B, Martin C. 2001. An essential role for phoP in Mycobacterium tuberculosis virulence. Mol Microbiol 41:179–87.

29. Cooper AM. 2014. Mouse model of tuberculosis. Cold Spring Harb Perspect Med 5:a018556.

30. Capinos Scherer CF, Endsley JJ, de Aguiar JB, Jacobs WR, Jr., Larsen MH, Palmer MV, Nonnecke BJ, Ray Waters W, Mark Estes D. 2009. Evaluation of granulysin and perforin as candidate biomarkers for protection following vaccination with Mycobacterium bovis BCG or M. bovisDeltaRD1. Transbound Emerg Dis 56:228–39.

31. Akter S, Chauhan KS, Dunlap MD, Choreno-Parra JA, Lu L, Esaulova E, Zuniga J, Artyomov MN, Kaushal D, Khader SA. 2022. Mycobacterium tuberculosis infection drives a type I IFN signature in lung lymphocytes. Cell Rep 39:110983.

32. Andersen P, Scriba TJ. 2019. Moving tuberculosis vaccines from theory to practice. Nat Rev Immunol doi:10.1038/s41577-019-0174-z.

33. Kang DD, Lin Y, Moreno JR, Randall TD, Khader SA. 2011. Profiling early lung immune responses in the mouse model of tuberculosis. PLoS One 6:e16161.

34. Palmer MV, Thacker TC, Waters WR. 2007. Vaccination of white-tailed deer (Odocoileus virginianus) with Mycobacterium bovis bacillus Calmette Guerin. Vaccine 25:6589–97.

35. Palmer MV, Thacker TC, Rabideau MM, Jones GJ, Kanipe C, Vordermeier HM, Ray Waters W. 2019. Biomarkers of cell-mediated immunity to bovine tuberculosis. Vet Immunol Immunopathol 220:109988.

36. Waters WR, Palmer MV, Buddle BM, Vordermeier HM. 2012. Bovine tuberculosis vaccine research: historical perspectives and recent advances. Vaccine 30:2611–22.

37. Palmer MV, Thacker TC, Waters WR, Robbe-Austerman S, Aldwell FE. 2014. Persistence of Mycobacterium bovis bacillus Calmette-Guerin (BCG) Danish in white-tailed deer (Odocoileus virginianus) vaccinated with a lipid-formulated oral vaccine. Transbound Emerg Dis 61:266–72.

38. Waters WR, Maggioli MF, McGill JL, Lyashchenko KP, Palmer MV. 2014. Relevance of bovine tuberculosis research to the understanding of human disease: historical perspectives, approaches, and immunologic mechanisms. Vet Immunol Immunopathol 159:113–32.

39. Flynn JL, Gideon HP, Mattila JT, Lin PL. 2015. Immunology studies in non-human primate models of tuberculosis. Immunol Rev 264:60–73.

40. Larsen SE, Williams BD, Rais M, Coler RN, Baldwin SL. 2022. It Takes a Village: The Multifaceted Immune Response to Mycobacterium tuberculosis Infection and Vaccine-Induced Immunity. Front Immunol 13:840225.

41. Davis WC, Mahmoud AH, Abdellrazeq GS, Elnaggar MM, Dahl JL, Hulubei V, Fry LM. 2022. Ex vivo Platforms to Study the Primary and Recall Immune Responses to Intracellular Mycobacterial Pathogens and Peptide-Based Vaccines. Front Vet Sci 9:878347.

42. Abdellrazeq GS, Fry LM, Elnaggar MM, Bannantine JP, Schneider DA, Chamberlin WM, Mahmoud AHA, Park KT, Hulubei V, Davis WC. 2020. Simultaneous cognate epitope recognition by bovine CD4 and CD8 T cells is essential for primary expansion of antigen-specific cytotoxic T-cells following ex vivo stimulation with a candidate Mycobacterium avium subsp. paratuberculosis peptide vaccine. Vaccine 38:2016–2025.

43. Flynn JL, Chan J. 2022. Immune cell interactions in tuberculosis. Cell 185:4682–4702.

44. Gideon HP, Phuah J, Myers AJ, Bryson BD, Rodgers MA, Coleman MT, Maiello P, Rutledge T, Marino S, Fortune SM, Kirschner DE, Lin PL, Flynn JL. 2015. Variability in tuberculosis granuloma T cell responses exists, but a balance of pro- and anti-inflammatory cytokines is associated with sterilization. PLoS Pathog 11:e1004603.

45. Huang L, Nazarova EV, Tan S, Liu Y, Russell DG. 2018. Growth of Mycobacterium tuberculosis in vivo segregates with host macrophage metabolism and ontogeny. J Exp Med 215:1135–1152.

46. Grant NL, Maiello P, Klein E, Lin PL, Borish HJ, Tomko J, Frye LJ, White AG, Kirschner DE, Mattila JT, Flynn JL. 2022. T cell transcription factor expression evolves over time in granulomas from Mycobacterium tuberculosis-infected cynomolgus macaques. Cell Rep 39:110826.

47. McCaffrey EF, Donato M, Keren L, Chen Z, Delmastro A, Fitzpatrick MB, Gupta S, Greenwald NF, Baranski A, Graf W, Kumar R, Bosse M, Fullaway CC, Ramdial PK, Forgo E, Jojic V, Van Valen D, Mehra S, Khader SA, Bendall SC, van de Rijn M, Kalman D, Kaushal D, Hunter RL, Banaei N, Steyn AJC, Khatri P, Angelo M. 2022. The immunoregulatory landscape of human tuberculosis granulomas. Nat Immunol 23:318–329.

48. Nelp MT, Kates PA, Hunt JT, Newitt JA, Balog A, Maley D, Zhu X, Abell L, Allentoff A, Borzilleri R, Lewis HA, Lin Z, Seitz SP, Yan C, Groves JT. 2018. Immune-modulating enzyme indoleamine 2,3-dioxygenase is effectively inhibited by targeting its apo-form. Proc Natl Acad Sci U S A 115:3249–3254.

49. Sun C, Mezzadra R, Schumacher TN. 2018. Regulation and Function of the PD-L1 Checkpoint. Immunity 48:434–452.

50. Kanipe C, Boggiatto PM, Putz EJ, Palmer MV. 2022. Histopathologic differences in granulomas of Mycobacterium bovis bacille Calmette Guerin (BCG) vaccinated and non-vaccinated cattle with bovine tuberculosis. Front Microbiol 13:1048648.

51. Palmer MV, Thacker TC, Kanipe C, Boggiatto PM. 2021. Heterogeneity of Pulmonary Granulomas in Cattle Experimentally Infected With Mycobacterium bovis. Front Vet Sci 8:671460.

52. Atkinson GC, Tenson T, Hauryliuk V. 2011. The RelA/SpoT homolog (RSH) superfamily: distribution and functional evolution of ppGpp synthetases and hydrolases across the tree of life. PLoS One 6:e23479.

53. Dalebroux ZD, Svensson SL, Gaynor EC, Swanson MS. 2010. ppGpp conjures bacterial virulence. Microbiol Mol Biol Rev 74:171–99.

54. Kundra S, Colomer-Winter C, Lemos JA. 2020. Survival of the Fittest: The Relationship of (p)ppGpp With Bacterial Virulence. Front Microbiol 11:601417.

55. Dozot M, Boigegrain RA, Delrue RM, Hallez R, Ouahrani-Bettache S, Danese I, Letesson JJ, De Bolle X, Kohler S. 2006. The stringent response mediator Rsh is required for Brucella melitensis and Brucella suis virulence, and for expression of the type IV secretion system virB. Cell Microbiol 8:1791–802.

56. Charity JC, Blalock LT, Costante-Hamm MM, Kasper DL, Dove SL. 2009. Small molecule control of virulence gene expression in Francisella tularensis. PLoS Pathog 5:e1000641.

57. Bugrysheva JV, Godfrey HP, Schwartz I, Cabello FC. 2011. Patterns and regulation of ribosomal RNA transcription in Borrelia burgdorferi. BMC Microbiol 11:17.

58. Taylor CM, Beresford M, Epton HA, Sigee DC, Shama G, Andrew PW, Roberts IS. 2002. Listeria monocytogenes relA and hpt mutants are impaired in surface-attached growth and virulence. J Bacteriol 184:621–8.

59. Park S-I, Jeong J-H, Choy HE, Rhee JH, Na H-S, Lee T-H, Her M, Cho K-O, Hong Y. 2010. Immune response induced by ppGpp-defective Salmonella enterica serovar Gallinarum in chickens. The Journal of Microbiology 48:674–681.

60. Pizarro-Cerda J, Tedin K. 2004. The bacterial signal molecule, ppGpp, regulates Salmonella virulence gene expression. Mol Microbiol 52:1827–44.

61. Geiger T, Kastle B, Gratani FL, Goerke C, Wolz C. 2014. Two small (p)ppGpp synthases in Staphylococcus aureus mediate tolerance against cell envelope stress conditions. J Bacteriol 196:894–902.

62. Davis WC, Abdellrazeq GS, Mahmoud AH, Park KT, Elnaggar MM, Donofrio G, Hulubei V, Fry LM. 2021. Advances in Understanding of the Immune Response to Mycobacterial Pathogens and Vaccines through Use of Cattle and Mycobacterium avium subsp. paratuberculosis as a Prototypic Mycobacterial Pathogen. Vaccines (Basel) 9.

